# The Use and Misuse of Composite Environmental Indices

**DOI:** 10.1101/2022.03.15.484501

**Authors:** Shelley M. Fischer, Michael K. Joy, Wokje Abrahamse, Taciano L. Milfont, Lynda M. Petherick

## Abstract

Composite indices have been widely used to rank the environmental performance of nations. Such environmental indices can be useful in communicating complex information as a single value and have the potential to generate political and media awareness of environmental issues. However, indices that are poorly constructed can hinder efforts to identify environmental failings. Here, we provide a critical review of the theoretical and methodological foundations of environmental indices to enhance our understanding of the accuracy and applicability of such indices. In the present study we classify existing indices according to ranking goal, measurement components, and weighting methods. Using New Zealand and Niger as case studies, we examine correlations between ranks in ten national level indices to outline how measurement components and the goal of ranking itself may provide a more, or less, optimistic view of the state of a country’s environment. Our results suggest that environmental indices that include human health, socioeconomic, and policy indicators (such as human access to sanitation and clean drinking water) are positively correlated with each other, and that those excluding human health, socioeconomic, and policy indicators are also positively correlated with each other, while these two types of environmental indices are negatively correlated to each other. Our results demonstrate that the inclusion of indicators that do not relate to the actual state of natural environments can confound results. When choosing an existing environmental index – or developing a new one – it is important to assess whether the ranking goal and the included indicators are appropriate. This is important because the inclusion of confounding indicators in environmental indices may misrepresent the actual state of natural environments.

## Introduction

Composite environmental indices attempt to communicate complex information by combining multiple measures into a single indicator. They provide an overall snapshot of some features of environmental systems and allow for comparisons between environmental systems (1). Environmental indices can be useful tools to measure stability and change over time (improvement or decline), easily communicate information to the general public and policy makers, drive accountability between countries, encourage accountability of a nation’s government to its citizens, and facilitate public engagement (2, 3). Environmental indices can also help inform and support policy decisions, as well as generate political and media awareness of environmental issues (4). However, if indices are poorly constructed they can hinder efforts to identify and address environmental failings, and may mislead policy messages and decisions (2, 5). To enhance our understanding of the accuracy and applicability of a given environmental index, it is crucial that the underlying theories, ranking goals, and method(s) used to develop the index are considered.

Although the same basic steps (i.e., development of a theoretical framework, data selection, imputation of missing data, multivariate analysis and normalisation, weighting and aggregation, uncertainty and sensitivity analysis) are typically used to develop composite indices, existing indices can differ widely in terms of their geographic scope, ranking goals, measurement components, terminology, and statistical methods (2, 3). Hence, several issues should be considered when assessing the quality of an environmental index and whether it is fit for purpose (6, 7), as summarised in Table 1. Firstly, environmental indices differ according to the theoretical frameworks that inform them. This is reflected in the ranking goals and measurement components that researchers adopt. Some indices, such as the Environmental Performance Index (EPI) and Composite Index of Environmental Performance (CIEP) focus on sustainability as a whole and include one or more of the three interconnected dimensions of sustainability (environment, society, economy – see Fig 1). This is reflected in the ranking goals and the measurement components included in the index (7, 8). Environmental indices that reflect multiple dimensions of sustainability typically incorporate human health and socioeconomic indicators, as well as indicators of environmental policies and practices (either directly or as a calculation component). Other indices, such as the Environmental Vulnerability Index (EVI), Proportional Environmental Impact Rank (pENV), and Absolute Environmental Impact Rank (aENV), focus on the environmental dimension of sustainability and explicitly exclude human health and socioeconomic indicators to avoid confounding or diluting the environmental component (7, 9).

**Table 1.** Issues for Consideration When Assessing the Quality of an Environmental Index and Whether it is Fit for Purpose.

| Issue for consideration | Summary of problem |
| --- | --- |
| Theoretical framework | Indices have different ranking goals and measurement components based on the theoretical components targeted. Indices can be broadly split into two types, they either:<br>- reflect multiple dimensions of sustainability and include human health, socioeconomic, and policy indicators, or<br>- focus exclusively on the environmental pillar of sustainability and exclude human health, socioeconomic, and policy indicators. |
| Indicator choice | Some indices use:<br>- low number of indicators<br>- unsuitable indicators chosen<br>- exclude key indicators. |
| Data quality and choice | Some indices use:<br>- insufficient data<br>- data of dubious accuracy. |
| Terminology used | Variations in terminology can:<br>- lead to confusion about what is being measured (e.g., the name of the index does not reflect what is being measured)<br>- lead to confusion between indices (e.g., indices share the same/similar name). |
| Weighting and aggregation techniques | Different weighting and aggregation techniques (expert opinion, equal weighting, statistics-based weighting) are used and can impact rankings. |

**Figure 1.**
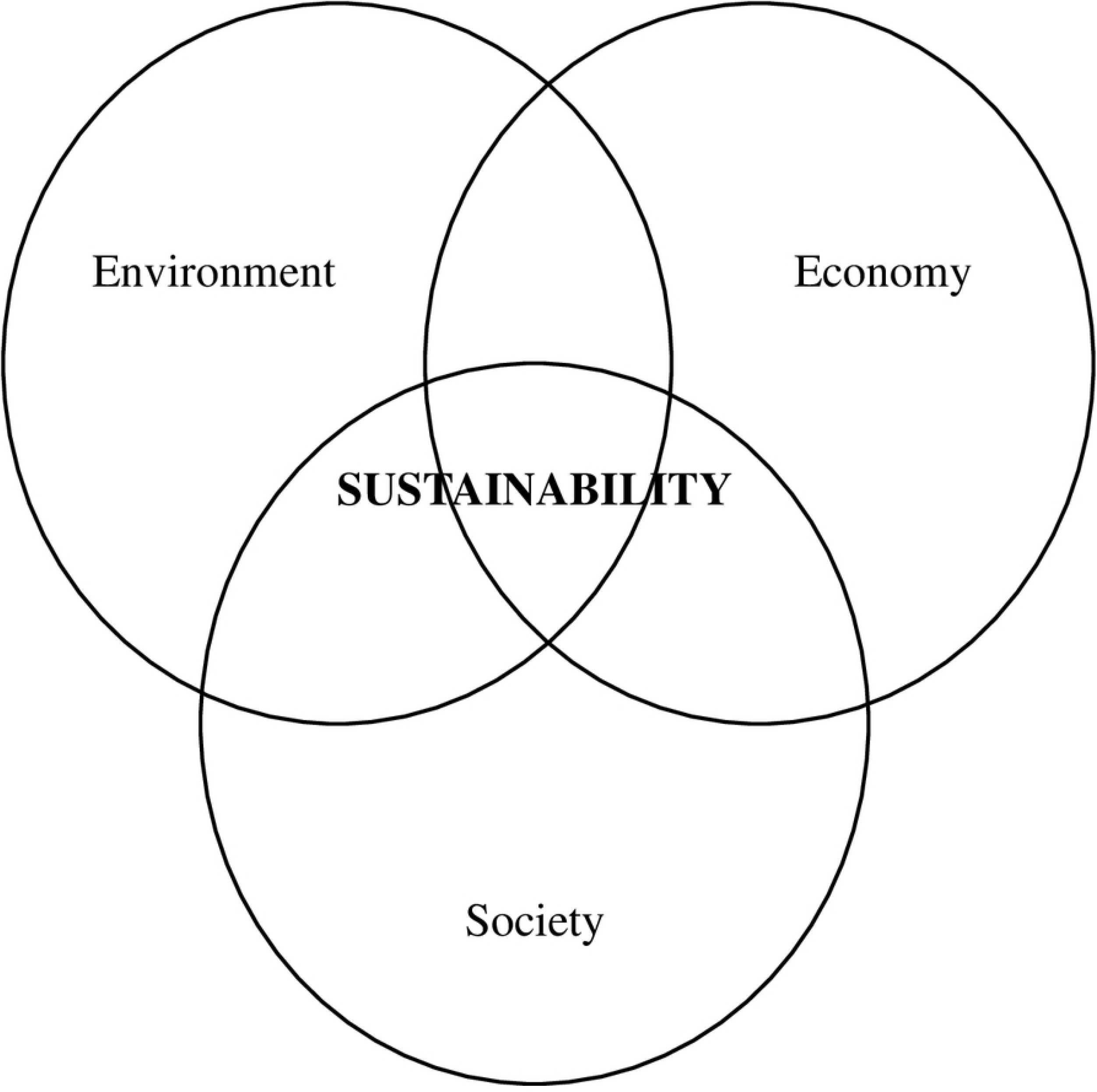
Venn Diagram Depicting the Three Overlapping Dimensions of Sustainability, Adapted from Brundtland (10).

Secondly, researchers have questioned the quality of certain indices because of indicator choice and the exclusion of core indicators. The EPI in particular has been criticised for using a small number of indicators and for excluding key indicators for water quality, contamination, erosion, and waste management. This may limit the ability of the index to capture and assess environmental quality (4, 11). It is worth noting that there is often a compromise between scientific precision and information available—what scientists would like to measure does not always correlate with what can practically be measured at a given point in time (3). However, in some cases, unsuitable indicators are chosen when more appropriate indicators are available. For example, the CIEP and EPI include human health indicators such as human access to sanitation and clean drinking water (12, 13) when the quality of sanitation and sewerage treatment would be more appropriate in assessing environmental state (11).

Thirdly, data quality and choice are a critical problem for many environmental indices. For example, the EPI includes a human health indicator: the number of age standardised disability adjusted life years lost per 100,000 persons due to exposure to fine air particulate smaller than 2.5μm (PM_2.5_). This is a problematic choice because (1) it is primarily a human health indicator, and (2) the quality of the data is questionable. Because direct modelling of PM_2.5_ is still rare in most parts of the world, modelled data are used instead (14). Modelled data should be treated with caution because it is based on assumptions and limited data (15, 16). To illustrate, New Zealand is ranked 1^st^ equal for PM_2.5_ in the EPI, but direct monitoring for particulate matter in New Zealand is both temporally and spatially sporadic (17). From the limited data available, it seems unlikely that New Zealand would be a top performer given that 37% of sites monitored in New Zealand for the period 2014-2016 exceed the World Health Organisation’s (WHO) annual average PM_2.5_ guidelines for long-term exposure (18). Another example of questionable data quality and coverage is the inclusion of renewable water resources and freshwater withdrawal in the Legatum Natural Environment Index and Environmental State and Sustainability Index (ESSI) (1, 19). Because data is limited for large parts of the world, modelled data from the Food and Agriculture Organisation (FAO) AQUASTAT database is used instead. The FAO itself raised the need for caution when using the dataset: “the problems of insufficient data and data of dubious accuracy paint a picture that, if interpreted as a final product, lead to incorrect assumptions” (20, Challenges section, para. 5) The FAO highlighted issues with data coverage for North America, Europe, Japan, Australia, and New Zealand — all of which rank highly in the Legatum Natural Environment Index (21).

Fourthly, different terminology is used and this variation can lead to debate and confusion (11, 22). An example of this is the EPI, previously called the Environmental Sustainability Index (ESI). Despite the name change, it retains a sustainability focus while still incorporating human health and socioeconomic indicators. Furthermore, numerous environmental indices have the same name but contain different indicators and have different purposes. For example, Environmental Quality Index (EQI) in Argentina measures living conditions for city districts in Buenos Aires (23), while an EQI for 3141 counties in the United States assessed environmental conditions as a predictor of human health outcomes (24). While the two EQI share the same name, the scale and focus of the indices are quite different.

Finally, there are issues with the statistical methods used for some environmental indices. Different procedures may be used for imputing missing values, normalisation, weighting, and aggregation. Weighting techniques are particularly important as they can lead to different rankings (2, 25, 26). There are three common methods for weighting indicators: (1) expert-opinion based weighting, which uses subjective judgements in an open discussion process with experts (e.g., EPI), (2) equal weighting where all indicators are given the same weight (e.g., EVI), and (3) statistics-based weighting (e.g., ENV). Indices that use equal and expert opinion-based weighting methods have been criticised for failing to consider the interlinkages and dynamic interrelations of the various components (26–28).

To avoid such weighting problems, indices such as the ENV and ESSI use multivariate analysis to explore the nature of datasets and determine weights (1, 7). The ESSI, which was developed as an alternative to the EPI, illustrates how weighting techniques can impact on country rankings. While there are a few differences in variable choice, the dimensions of the EPI and ESSI are fairly similar (1). The ESSI uses factor analysis instead of the expert opinion-based weighting used for the EPI and the two indices show marked differences in the weights assigned to different indicators and dimensions. There are also pronounced differences in the rankings of some countries; for example, the United States is ranked 47/163 in the ESSI and 26/163 in the EPI (1). This example illustrates the impact of different weighting techniques and how important it is that developers of environmental indices explicitly and transparently communicate the statistical methods used.

Beyond these issues for assessing environmental indices, previous research has also highlighted inconsistencies and variations in how nations are ranked in existing environmental and sustainability indices, highlighting the need for further research. Bradshaw et al. (2010) compared country ranks in the pENV with rankings in two environmental indices, the EPI and the Ecological Footprint (it is unclear which version of the Ecological Footprint was used – per person or total). These authors found weak negative correlations between the pENV and EPI (Kendall’s τ = −0.21, P = 0.0001, n = 149 countries) and a weak positive relationship between the pENV and Ecological Footprint (Kendall’s τ = 0.09, P = 0.0991, n = 150 countries). This suggests that there may be a higher degree of variation in country ranks between those environmental indices that include human health, socioeconomic, and policy indicators and those indices that exclude such indicators. In the same study, country ranks in the pENV, EPI, and Ecological Footprint were compared with country ranks in two other indices, the Genuine Savings Index (GSI) and the Human Development Index (HDI). The GSI measures economic sustainability and the HDI measures life expectancy, education, and per capita income. A moderate to strong correlation was observed between the EPI and HDI (Kendall’s τ = 0.70, P = 0.0001, 110 countries), a moderate negative correlation between the Ecological Footprint and the HDI (Kendall’s τ = −0.67, P = 0.0001, 110 countries), and weak negative relationships between the pENV and EPI (Kendall’s τ = −0.21, P = 0.0001, 149 countries), pENV and HDI (Kendall’s τ = −0.22, P < 0.0001, 178 countries, and pENV and GSI (Kendall’s τ = −0.25, P = 0.0001, 118 countries). These results indicate that indicator choice may play a role in rank variation and that ranks in environmental indices such as the EPI—which includes human health, socioeconomic, and policy indicators—are similar to rankings in general sustainability and economic indices. However, Bradshaw et al. (2010) used only four environmental indices: their own aENV and pENV, the EPI, and Ecological Footprint. We have identified six further environmental indices for which country rankings can be compared to improve our knowledge of how ranking goals, measurement components, and weighting methods may influence rank variation.

In this study, we advance the work of Bradshaw et al. (2010) by comparing country rankings across all 10 environmental indices we are aware of that include a minimum of 100 countries: the CIEP, Ecological Footprint (per person and total), EPI, ENV (absolute and proportional), ESSI, EVI, Ecosystem Wellbeing Index (EWI), and Legatum Natural Environment. Specifically, we: (1) summarise existing indices according to ranking goal, measurement components, and weighting methods, (2) test for correlations between ranks in existing national level indices, and (3) use New Zealand and Niger as case studies, outline how measurement components and the goal of ranking itself can influence the ranking of nations. New Zealand and Niger were chosen because in our preliminary analysis we noted that the two countries commonly featured among the best or worst performers. For example, in the 2020 version of the EPI New Zealand ranks at 19 and Niger ranks at 152. In contrast, New Zealand ranks at 161 in the pENV and Niger ranks at 5. New Zealand and Niger represent two contrasting country profiles that illustrate how indices which include human health and socioeconomic indicators may offer an overly optimistic (or pessimistic) view of a country’s environmental performance according to its development status. New Zealand provides an example of a developed nation with a high standard of living as measured by the HDI (29), with widespread access to sanitation and clean drinking water (30, 31). Environmental issues in New Zealand include water pollution, soil degradation, high greenhouse gas emissions per person, and biodiversity loss (32). In contrast, Niger is a developing country that consistently ranks near the bottom of the HDI (33). In Niger, air pollution from cooking over open flames is common and access to clean drinking water and sanitation is poor (34–36). Environmental issues in Niger include land degradation and desertification, pressure on natural resources from population growth, and pollution from mining (37, 38).

Our study aims to quantify inconsistencies and wide variations in ranking amongst existing environmental indices and establish the impact of indicator choice on global ranks. We demonstrate that the inclusion of human health, socioeconomic, and policy indicators confounds results providing a misleading view of a country’s environmental performance. In doing so, this paper provides a novel contribution to the development of composite environmental indices. Existing indices which include human health, socioeconomic and policy indicators should be used with caution. When developing a new environmental index human health, socioeconomic, and policy indicators should be explicitly excluded to accurately represent the true state of environments.

## Methods

After performing a survey of worldwide environmental indices we identified 12 potential national level environmental indices for inclusion in our study. Indices were excluded from our analysis if there was insufficient information about the indicators included, the weighting and aggregation methods, or there were less than 100 countries included. Under these criteria two indices were excluded, the Environmental Pressure Index (39), and the Environmental Degradation Index (27). We then examined the rank correlations for 10 national level environmental indices: the CIEP, Ecological Footprint (per person and total), EPI, ENV (absolute and proportional), ESSI, EVI, Ecosystem Wellbeing Index (EWI), and Legatum Natural Environment. Data for the Ecological Footprint, EPI and Legatum Natural Environment was sourced from official websites (40–42). Data for ENV, EVI, and ESSI was obtained from academic publications (1, 7, 9). Data for the CIEP was obtained directly from Dr Thiago Almeida (Campina Grande Federal University, Brazil), one of the developers of the CIEP (43).

To determine the degree of agreement between ranks we used Kendall’s τ coefficient, which is a commonly used measure of rank correlation (44). First, ranks were compiled for each index. Countries were only included if there was a value for each index, resulting in a total of 114 countries. Original rankings were normalised and Kendall’s τ was calculated using IBM SPSS Statistics (Statistical Package for Social Sciences) for Windows, Version 21.0.

We also investigated rank correlations for indices which were published in the same, or consecutive years. Original ranks were normalised and Kendall’s τ coefficient was used to determine the degree of agreement between ranks for: the pENV (2010) and EPI (2010), the aENV (2010) and EPI (2010), the Ecological Footprint total (2017) and EPI (2018), and the Ecological Footprint per person (2017) and EPI (2018).

The 10 national-level indices are summarised and grouped according to their aggregation and weighting method, and whether they include human health, socioeconomic, and policy indicators. For our two country case studies (New Zealand and Niger), we compiled a summary list of ranks in the CIEP, Ecological Footprint (per person and total), EPI, ENV (absolute and proportional), ESSI, EVI, EWI, and Legatum Natural Environment. Ranks were normalised 0-100.

## Results

### Summary of Existing Indices

We reviewed the ranking goals, measurement components, and weighting and aggregation methods of 10 existing composite environmental indices (Table 2). Four indices (the CIEP, EPI, ESSI, and Legatum Natural Environment) focus on environment impacts for human health or prosperity. These indices encompass multiple aspects of sustainability and include human health, socioeconomic, and policy indicators. The remaining six indices (both versions of the Ecological Footprint, the EVI, EQI, aENV, and pENV) focus on ecosystem health and environmental impacts. These indices focus solely on the environmental component of sustainability and explicitly exclude human health, socioeconomic, and policy indicators.

**Table 2.** Summary of Ten National-Level Environmental Indices Outlining Goal of Ranking Measurement Components, Weighting and Aggregation.

| Index name and source | Countries or regions included | Indicators included | Weighting and aggregation choices | Human health, socioeconomic, or policy indicators? | Goal of Ranking (ranked according to?) | First published in |
| --- | --- | --- | --- | --- | --- | --- |
| CIEP | 152 | 19 | Statistics based – CRITIC (Criteria Importance Through Intercriteria Correlation) | √ | Environmental performance (effects on human health) | Almeida, 2015 |
| Ecological Footprint (per person and total) | 178 countries + 9 OT | 16 over 6 issue categories | Accounting system (supply and demand of nature). Aggregation is expert opinion based (under development by JRC) | × | Ecological deficit/ecological reserves for a given human population | Rees, 1992 |
| EPI | 180 | 32 over 11 issue categories & 2 policy objectives | Expert-opinion based. Statistical audits by Joint Research Commission (2006, 2006, 2008, 2010, 2012, 2014, 2020) | √ | Environmental performance – proximity to established international environmental policy goals and targets | Esty et al., 2006 |
| ESSI | 163 | 22 over 9 issue categories | Statistics based – factor analysis | √ | State of environment and environmental sustainability | Oțoiu & Grădinaru, 2018 |
| EVI | 192 countries + 44 OT | 50 divided into 3 sub-indices | Equal weighting (1:1:1...) | × | Environmental vulnerability | Kaly et al., 1999 |
| EWI | 180 | 52 over 5 issue categories | Unweighted averages for land, water, air, species and genes, and resources use. Or, averages of land, water, air, species and genes | × | Environmental quality – proximity to goal (low level of ecosystem stress) | Prescott-Allen, 2001 |
| aENV (absolute rank) | 171 | 12 | Statistics based – countries ranked for each indicator then tested for robustness to inclusion/exclusion of measurement components | × | Environmental impact – performance with respect to total resources available | Bradshaw et al., 2010 |
| pENV (proportional rank) | 178 + 1 OT | 12 | ” | × | Environmental impact – performance with respect to total resources available for a given human population | Bradshaw et al. 2010 |
| Legatum Natural Environment | 167 |  | Weights determined by two factors: (1) relevance & significance of indicator to prosperity (informed by expert opinion & academic literature), and, to a lesser degree (2) statistical significance of indicator to economic and social wellbeing of a country. | √ | Parts of the physical environment that have a direct effect on people in their lives, and changes that might impact prosperity of future generations. | Legatum Institute 2017 |

**Table 2.** Summary Results of Kendall’s τ Examining Rank Correlations for Ten National-Level Environmental Indices (*n* = 114 countries): the EPI (2020), ESSI (2018), CIEP (2015), EVI (1999), EWI (2001), Ecological Footprint per person and total (2017), aENV, and pENV (2010), Legatum Natural Environment (LNE) (2021).

|  |  |  |  |  |  |  |  |  |  |
| --- | --- | --- | --- | --- | --- | --- | --- | --- | --- |
| EPI | 0.573** |  |  |  |  |  |  |  |  |
| ESSI | 0.574** | 0.683** |  |  |  |  |  |  |  |
| LNE | 0.448** | 0.491** | 0.451** |  |  |  |  |  |  |
| aENV | 0.110 | -0.037 | -0.067 | -0.089 |  |  |  |  |  |
| pENV | -0.034 | -0.085 | -0.136* | -0.050 | 0.274** |  |  |  |  |
| EF pp | -0.496** | -0.541** | 0.429** | -0.366** | -0.020 | -0.021 |  |  |  |
| EF tot | 0.025 | -0.130* | -0.123 | -0.069 | 0.499** | 0.221** | 0.205** |  |  |
| EVI | -0.121 | -0.259** | -0.262** | -0.058 | 0.079 | 0.438** | 0.127* | 0.189** |  |
| EWI | -0.260** | -0.399** | -0.355** | -0.095 | 0.171** | 0.262** | 0.381** | 0.300** | 0.403** |
|  | CIEP | EPI | ESSI | LNE | aENV | pENV | EF pp | EF tot | EVI |

Degree of correlation
| moderate negative | weak negative | very weak negative | very weak positive | weak positive | moderate positive | strong positive |
| --- | --- | --- | --- | --- | --- | --- |
| -0.59 to -0.40 | -0.39 to -0.20 | -0.19 to -0.01 | 0.00 to 0.19 | 0.20 to 0.39 | 0.40 to 0.59 | 0.60 to 0.79 |

Note:
\*\* Correlation is significant at the 0.01 level (2-tailed).
\*Correlation is significant at the 0.05 level (2-tailed).

**Table 3.** Rankings of New Zealand and Niger According to 10 National-Level Environmental Indices.

| Index | New Zealand's rank | Re-scaled 0-100 | Niger's rank | Re-scaled 0-100 | Human health, socioeconomic, policy indicators included? | Weighting |
| --- | --- | --- | --- | --- | --- | --- |
| CIEP (2015) | 22/152 | 13.9 | 125/152 | 82.1 | ✓ | statistics based |
| EPI (2020) | 19/180 | 10.1 | 152/180 | 68.2 | ✓ | expert opinion based |
| ESSI (2018) | 35/164 | 21.0 | 157/164 | 95.7 | ✓ | statistics based |
| EF per person (2017) | 132/175 | 75.3 | 51/175 | 28.7 | × | accounting based |
| EF total (2017) | 84/175 | 44.7 | 114/175 | 64.9 | × | accounting based |
| Legatum Natural Environment (2020) | 4/167 | 1.8 | 128/126 | 76.5 | ✓ | expert opinion and statistics based |
| aENV (2010) | 125/171 | 72.9 | 26/171 | 14.7 | × | statistics based |
| pENV (2010) | 161/178 | 90.4 | 5/178 | 2.3 | × | statistics based |
| EVI (1999) | 89/192 | 46.1 | 7/192 | 3.1 | × | equal weighting |
| EWI (2001) | 113/179 | 62.9 | 40/179 | 21.9 | × | unweighted averages |

The weighting and aggregation methods used are mixed. To illustrate, the four indices which account for multiple aspects of sustainability use three different methods. The CIEP and the ESSI use statistics-based weighting, the EPI uses expert opinion-based weighting, and the Legatum Natural Environment uses a combination of the two methods where weights are primarily determined by expert opinion and academic literature, and to a lesser extent by statistical significance to the economic and social wellbeing of a country. Indices which focus solely on the environmental component of sustainability also use a mixture of different methods. The aENV, and pENV use statistics-based weighting, the Ecological Footprint uses expert opinion-based weighting, and the EVI and EWI use equal weighting.

### Rank Correlations in Existing National-Level Environmental Indices

If these distinct environmental indices assess similar measurable aspects of environmental features of countries, moderate-to-high positive correlations are to be expected. However, a range of correlations were found, from moderate negative to strong positive, as presented in Table 2. The highest positive correlation is between the EPI and ESSI (Kendall’s τ = 0.683, P < 0.01). This positive correlation indicates that the increase in a country’s rank in the EPI is associated with an increase in rank in the ESSI. There were moderate positive correlations between the CIEP and EPI (Kendall’s τ = 0.573, P < 0.01), CIEP and ESSI (Kendall’s τ = 0.574, P < 0.01), CIEP and Legatum Natural Environment (Kendall’s τ = 0.448, P< 0.01), ESSI and Legatum Natural Environment (Kendall’s τ = 0.451, P < 0.01), EPI and Legatum Natural Environment (Kendall’s τ = 0.491, P < 0.01), aENV and Ecological Footprint (total) (Kendall’s τ = 0.499, P < 0.01), EVI and EWI (Kendall’s τ = 0.403, P < 0.01), and EVI and pENV (Kendall’s τ = 0.438, P < 0.01). There are weak positive correlations between the Ecological Footprint (per person) and EWI (Kendall’s τ = 0.381, P < 0.01), Ecological Footprint (total) and EWI (Kendall’s τ = 0.300, P < 0.01), aENV and pENV (Kendall’s τ = 0.274, P < 0.01), EWI and pENV (Kendall’s τ = 0.262, P < 0.01), and Ecological Footprint (per person) and Ecological Footprint (total) (Kendall’s τ = 0.205, P < 0.01).

There is a moderate negative correlation between the EPI and Ecological Footprint (per person) (Kendall’s τ = −0.541, P < 0.01). This negative correlation indicates there is an inverse relationship between the two indices, i.e., the increase in a country’s rank in the Ecological Footprint is associated with a decrease in rank in the EPI (or vice versa). There are also moderate negative correlations between the Ecological Footprint (per person) and ESSI (Kendall’s τ = −0.429, P < 0.1), Ecological Footprint (per person) and CIEP (Kendall’s τ = − 0.496, P < 0.01). Weak negative correlations also exist between the EPI and EWI (Kendall’s τ = −0.399, P < 0.01), Ecological Footprint (per person) and Legatum Natural Environment (Kendall’s τ = −0.366, P < 0.01), EWI and ESSI (Kendall’s τ = −0.355), EVI and ESSI (Kendall’s τ = −0.262, P < 0.01), EWI and CIEP (Kendall’s τ = −0.260, P < 0.01), EVI and EPI (Kendall’s τ = −0.259, P < 0.01).

Moreover, there is a very weak negative correlations between ranks for the 2010 versions of the aENV and EPI (Kendall’s τ = −0.054, P < 0.001). Country ranks vary up to 86 places (normalised 0-100) (Fig 2a). There is also a very weak negative correlation between ranks for the 2010 versions of the pENV and EPI (Kendall’s τ = −0.156, P < 0.001). Country ranks vary up to 98 places (normalised 0-100) (Fig 2b).

**Figure 2.**
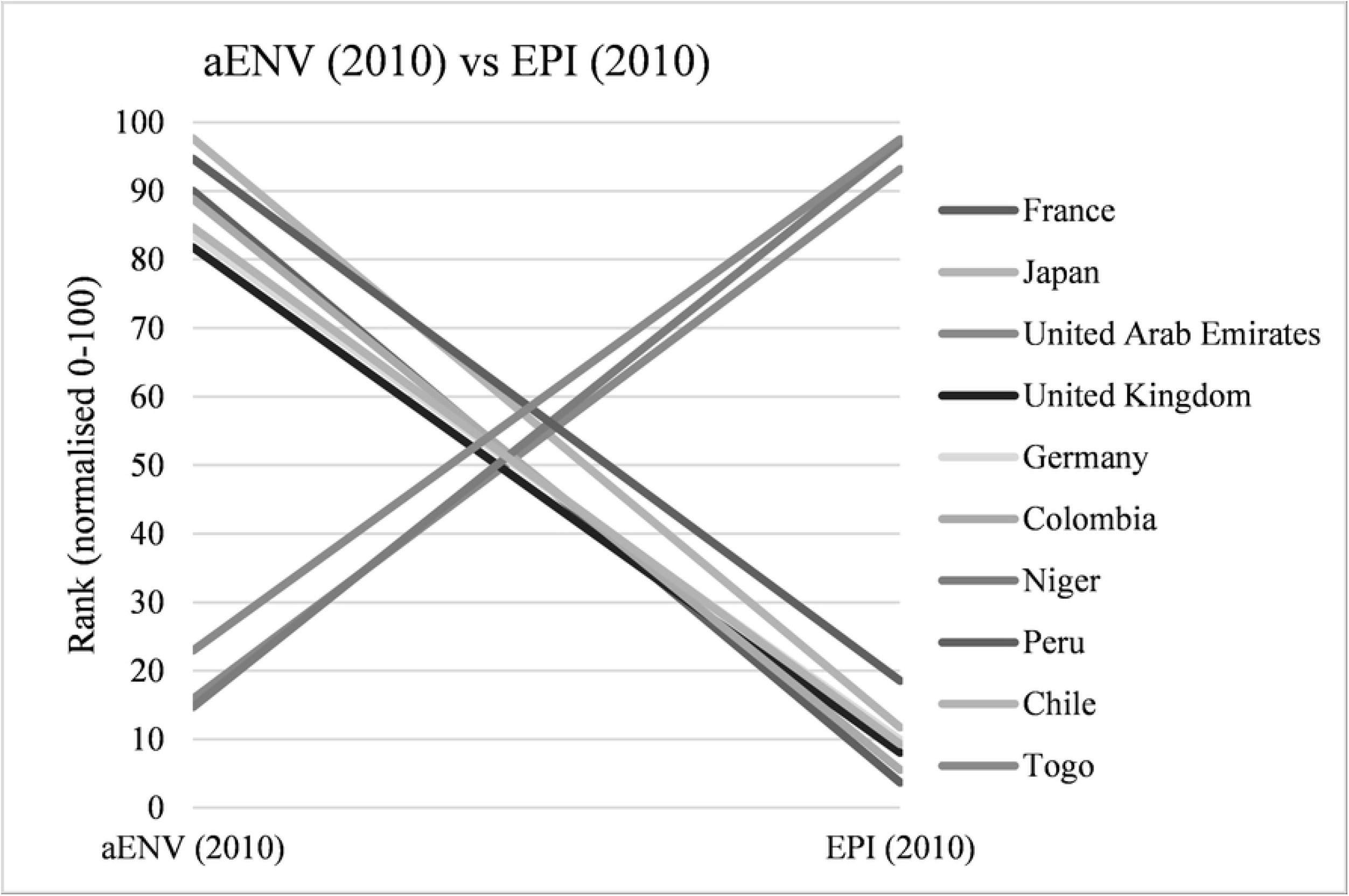
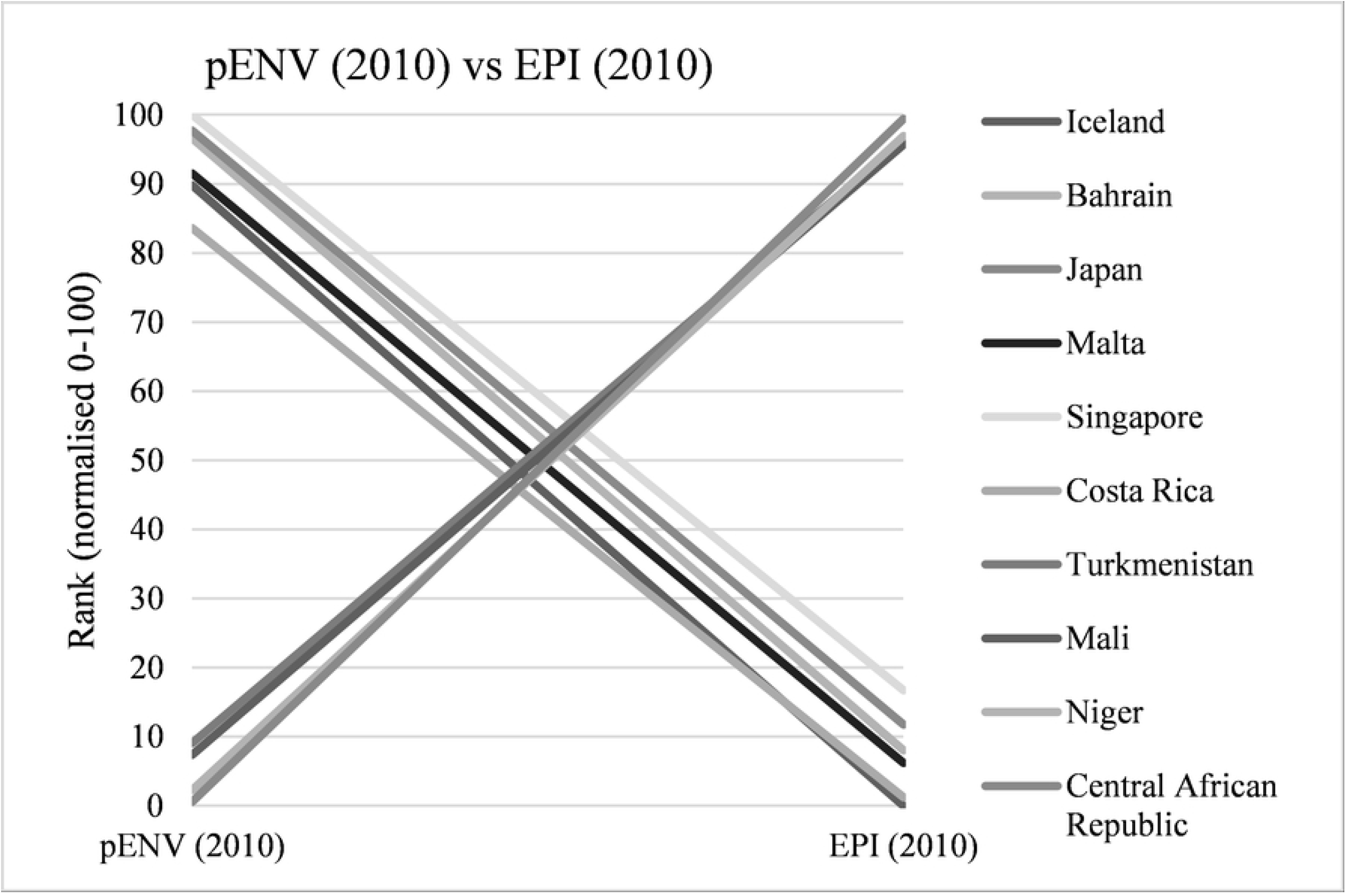
Countries with largest rank differences between the 2010 versions of the aENV, pENV and EPI. (A) 10 largest country rank differences between aENV and EPI (Kendall’s τ = −0.054, P < 0.01). (B) 10 largest country rank differences between pENV and EPI (Kendall’s τ = −0.156, P < 0.01).

There is a very weak negative correlation between ranks for the Ecological Footprint total (2017) and the EPI (2018) (Kendall’s τ = −0.125, P < 0.01). Ranks vary up to 94 places (normalised 0-100) (Fig 3a). There is a weak negative correlation between the Ecological Footprint per person (2017) and the EPI (2018) (Kendall’s τ = −0.206, P < 0.01). Ranks vary up to 95 places (normalised 0-100) (Fig 3b).

**Figure 3.**
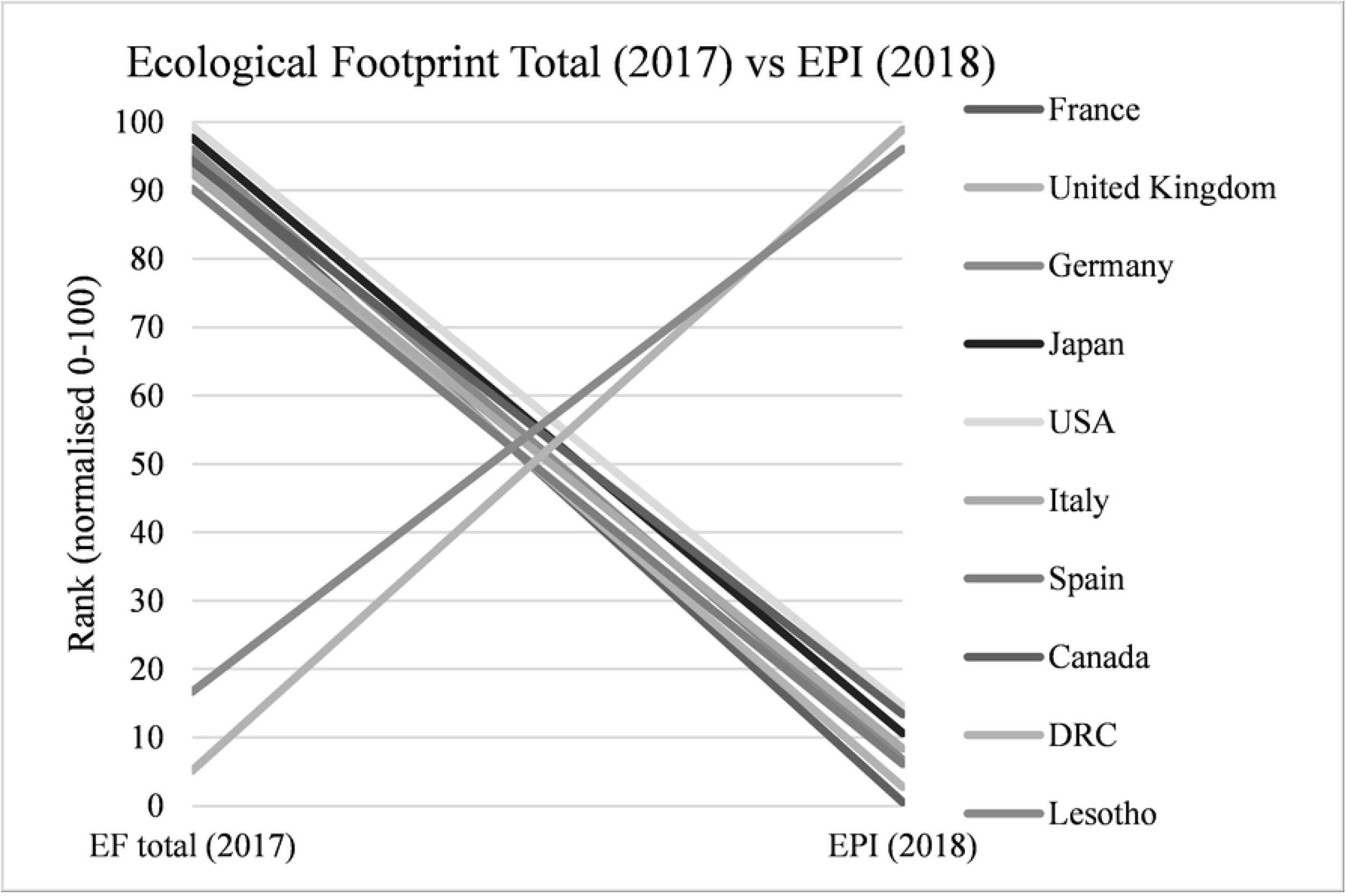
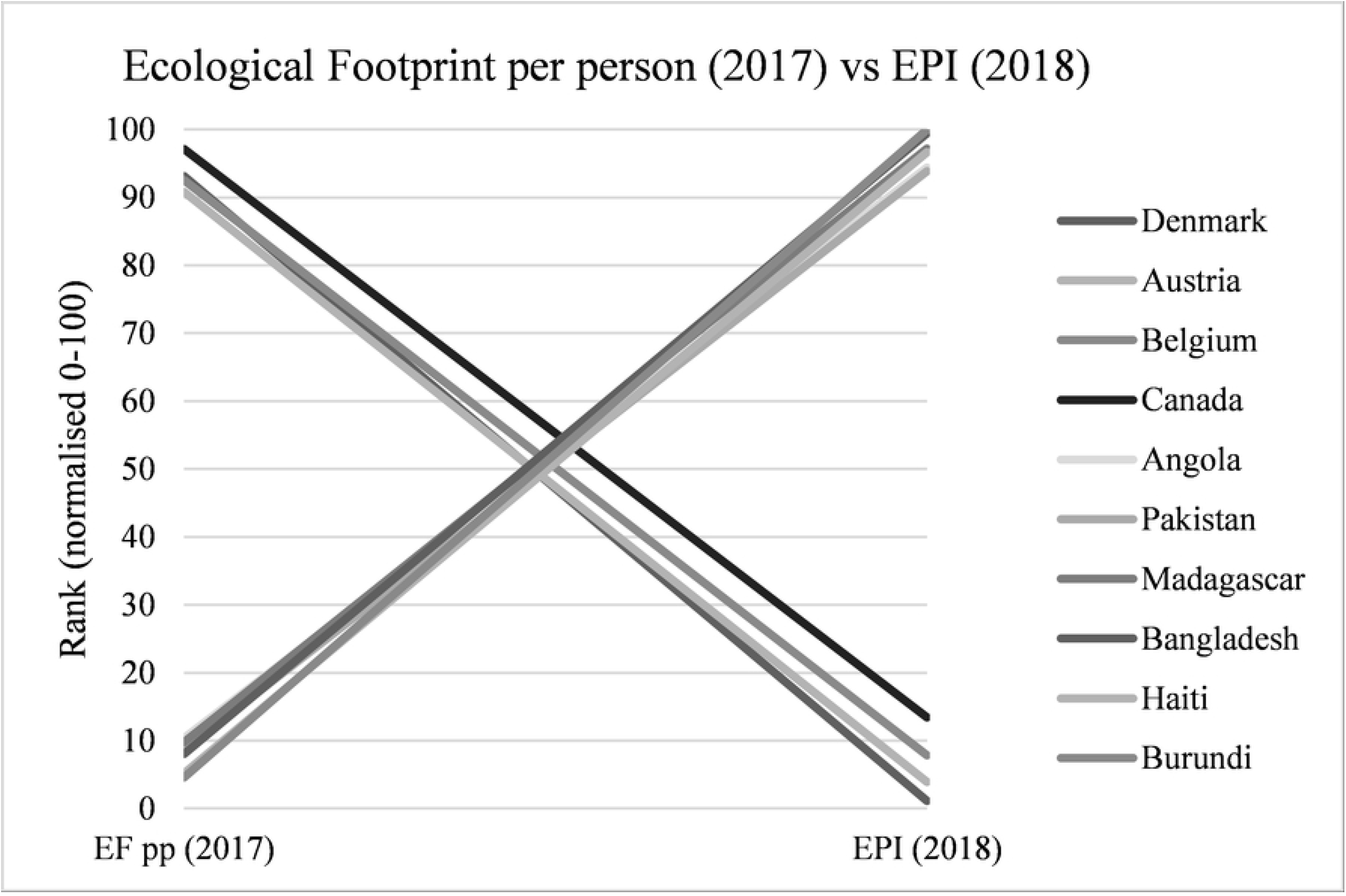
Countries with largest rank differences between the 2017/2018 versions of the Ecological Footprint (per person and total), and EPI. (A) 10 largest country rank differences between Ecological Footprint (total) and EPI (Kendall’s τ = −0.125, P < 0.01). Note. DRC = Democratic Republic of the Congo. (B) 10 largest country rank differences between Ecological Footprint (per person) and EPI.

### Case Studies: New Zealand and Niger

New Zealand and Niger are two countries which commonly feature among the best or worst performers across the different indices. New Zealand ranks between 19^th^ and 161^st^ across the different indices (Table 4). When normalised to a 0-100 scale New Zealand ranks between 10.1 and 90.4. New Zealand achieves the highest scores in indices that include human health, socioeconomic status, and policy indicators, while the lowest scores are in indices that exclude human health, socioeconomic, and policy indicators. Niger ranks between 5^th^ and 157^th^ across the different indices (Table 4). When normalised to a 0-100 scale New Zealand ranks between 2.3 and 57.7. Niger achieves the highest scores in indices that exclude human health, socioeconomic status, and policy indicators. Niger’s lowest scores are in indices that include human health, socioeconomic, and policy indicators.

The wide range in ranks for these two countries illustrates how indices which include human health, socioeconomic, or policy indicators may provide an overly optimistic (or pessimistic) view of the state of a countries environment according to the nations development status.

## Discussion

Composite indices have been widely used to rank the environmental performance of nations. Despite their benefits, previous research has posed critics to their theoretical or methodological foundations (see (4, 7, 8, 26). The aim of this paper is to quantify inconsistencies and wide variations in rank among existing environmental indices. Here we build upon the work of Bradshaw et al. (2010), demonstrating that the inclusion of human health, socioeconomic, and policy indicators may confound results, providing a misleading view of a country’s environmental state. Our results, based on correlations between rankings, indicate the following patterns: (1) indices that include human health, socioeconomic, and policy indicators are positively correlated with each other (CIEP, EPI, ESSI, Legatum Natural Environment), (2) indices that exclude human health, socioeconomic, and policy indicators are positively correlated with each other (per person and total Ecological Footprint, aENV, pENV, EVI, EWI), and (3) indices that include human health, socioeconomic, and policy indicators (CIEP, EPI, ESSI, Legatum Natural Environment) are negatively correlated with indices that exclude such indicators (per person and total Ecological Footprint, aENV, pENV, EVI, EWI).

While some correlations are weak, these general patterns indicate that ranking goal and indicator choice may influence a country’s rank among different indices. If the goal of ranking is to account for multiple dimensions of sustainability, then the inclusion of human health, socioeconomic, and policy indicators (either directly or as a calculation component) may be appropriate. However, if the goal of ranking is to account only for the environmental component of sustainability, then human health, socioeconomic, and policy indicators should not be included, as they have the potential to confound or dilute the environmental component of sustainability. Therefore, the actual state of natural environments may not be accurately represented.

One limitation in our correlation study is that the release dates for the indices range over several years. Unfortunately, some indices only have one iteration and it was therefore not possible to conduct a year-on-year comparison for every index. While this is an important limitation, analyses of indices such as the EPI have confirmed strong positive correlations between ranks over multiple iterations (see Table S1). This indicates that country ranks within the same index do not vary markedly over time, and provides some assurance in the results reported here.

Ranks for our two case studies (New Zealand and Niger) vary widely across different environmental indices. Niger scores well in indices that exclude human health, socioeconomic, and policy indicators. Niger is positioned 189/189 in the HDI (33) with 42.9% of the population living in extreme poverty (38). Only 13% of the population has access to basic sanitation services (34) and 23% of disability-adjusted life years (DALYs) can be attributed to lack of clean water, sanitation, and hygiene (35). Therefore, the inclusion of such indicators provides a less optimistic view of Niger’s environmental performance when compared to more developed nations.

By contrast, New Zealand scores well in indices that include human health, socioeconomic, and policy indicators. New Zealand is a developed country with a high standard of living and ranks 14/189 in the HDI (45). This is reflected in indicators such as access to clean drinking water, access to sanitation, and indoor air pollution (30). The inclusion of such indicators provides a more optimistic view of New Zealand’s performance when compared to less developed nations. New Zealand scores poorly in indices which include indicators for agricultural production and water quality and exclude human health, socioeconomic, and policy indicators. Primary production is a significant earner for New Zealand, with dairy products accounting for 42% of primary industry revenue in 2020 (46). New Zealand has weak agricultural policies, which are poorly implemented and environmental pollution from agriculture is significant (47–50). Environmental compliance, monitoring, and enforcement in New Zealand is poor and political interference in environmental management is commonplace (51–53). Therefore, the inclusion (or exclusion) of indicators for agriculture related pollution may influence New Zealand’s rankings in different indices. To illustrate, New Zealand scores poorly in both the aENV and pENV, which include fertiliser use and water pollution, but ranks well in the ESSI and CIEP, which do not account for agricultural pollution (1, 7, 13).

The wide variation in New Zealand’s ranking within different environmental indices demonstrates how indicator choice can lead to misleadingly high rankings, with the potential to misrepresent the actual state of natural environments. In turn, misleading rankings can be misinterpreted or misused by policy makers, the public, and stakeholders in different industries. As an example, the director of two industry organisations representing dairy farmers in New Zealand has used the county’s high ranking in the EPI to argue that the benefit of some environmental regulation in New Zealand is not clear because food is produced “more sustainably than other countries…in a natural environment in which we ranked high globally” (54, parag. 26). The same director has also used New Zealand’s high ranking in the Legatum Prosperity index to argue that New Zealand’s agricultural industry has a lower environmental impact than other countries (55).

Moreover, researchers have used a variety of weighting methods and there is no clear pattern across the different indices. For example, New Zealand and Niger have quite disparate rankings in the ESSI and pENV (20.1 and 90.4; 95.7 and 2.3, respectively), even though both indices use statistics-based weighting procedures. Consequently, while weighting procedures may have some impact on overall ranking, it is likely that indicator choice has a larger effect.

## Conclusions

In this article we have: (1) classified existing indices according to ranking goal, measurement components, and weighting methods, (2) tested for correlations between ranks in existing national level indices, (3) used Niger and New Zealand as case studies to outline how measurement components and the goal of ranking itself can influence the ranking of nations. Our findings highlight inconsistencies and wide variations in country ranks among existing environmental indices. Specifically, our results suggest that indicator choice can impact ranking, with the potential to misrepresent the actual state of environments. Using New Zealand and Niger as examples, we have demonstrated that the inclusion of human health indicators (such as access to clean drinking water and access to sanitation) may provide a more, or less, optimistic view of the state of a country’s environment. Indices that include specific dimensions of typical environmental problems in industrialised countries such as New Zealand (e.g., pollution from agricultural production, soil or waste management) may more accurately identify environmental failings.

The national-level rank comparison conducted in this study demonstrates the importance of ranking goal and measurement components when developing an environmental index. When choosing an existing environmental index, or developing a new one, it is important to assess whether the goal of ranking itself, and the indicators included are appropriate. Do they accurately represent the environmental system being assessed? Are construction processes open and transparent? Do human health, socioeconomic, or policy indicators confound or dilute the environmental component? Does the name of the index accurately reflect what it is measuring? Understanding the issues and challenges with constructing environmental indices means it is easier to choose or design an improved index. It is especially important to be clear about the ranking goals and measurement components of environmental indices when using them in combination with other indices to avoid confounding results.

Here we provide a novel contribution by quantifying inconsistencies and wide variations in rank among 10 existing environmental indices. Our correlation analysis illustrates how indicator choice can impact ranks with the potential to misrepresent the actual state of natural environments. Using the case studies of New Zealand and Niger, we have demonstrated how misleading rankings can be misinterpreted or misused by stakeholders in different industries. To avoid confounding results, environmental indices should focus solely on the environmental domain and explicitly exclude human health, socioeconomic, and policy indicators. This research will guide and support decision making for the development of new environmental indices, such as the New Zealand Environmental Quality Index (NZ— EQI) which is currently under development.

